# Mena (ENAH) Promotes KRAS-Driven Tumor Growth and Metastatic Progression in Pancreatic Ductal Adenocarcinoma

**DOI:** 10.64898/2026.01.07.698161

**Authors:** Lina A. Ariyan, Erika P. Zambalde, Jiufeng Li, Tanis T. McAuliffe, Elizabeth Ma, Francisco P. Martinez, Nicole C. Panarelli, Robert J. Eddy, Priyanka Patil, John Condeelis, Hava Gil-Henn, John C. McAuliffe

## Abstract

Metastatic pancreatic ductal adenocarcinoma (PDAC) remains incurable and is projected to become the second leading cause of cancer-related death by 2030. Despite therapeutic advances, median survival remains under one year. Activating KRAS mutations drive the majority of PDAC and underlie their highly aggressive behavior. Although emerging KRAS inhibitors show promise, resistance limits their clinical efficacy, underscoring the need to identify additional regulators of mutant *KRAS* signaling. We investigated the role of the actin-regulatory protein Mena (ENAH) in KRAS^G12D^-driven PDAC using novel *in vitro* and orthotopic *in vivo* models with Mena overexpression and knockdown. Mena overexpression markedly increased primary tumor growth and spontaneous liver metastasis, whereas Mena knockdown delayed tumor onset and reduced metastatic burden. Mechanistically, Mena depletion significantly decreased AKT and ERK activity downstream of KRAS in a SHIP2-mediated manner, indicating that Mena enhances oncogenic signaling required for PDAC progression. These findings reveal Mena as a critical modulator of KRAS^G12D^-driven pancreatic tumor growth and metastasis. By promoting key survival and proliferation pathways, Mena contributes to the aggressive phenotype characteristic of KRAS-mutant PDAC and represents a promising therapeutic target for patients with both locoregional and metastatic disease.

## INTRODUCTION

Pancreatic ductal adenocarcinoma (PDAC) is characterized by inevitable metastasis and resistance to therapy, making it the projected second leading cause of cancer-related death worldwide by 2030 ^1, 2^. Despite advances in multimodality management, survival outcomes remain dismal. Approximately 80% of patients present with metastatic disease ^1^, for whom combination cytotoxic chemotherapy provides a median overall survival of only ∼11 months ^3^. Even among patients eligible for curative-intent resection, 5-year survival rarely exceeds 25% and most ultimately succumb to metastatic progression ^4–6^. These observations underscore the critical need to define the biological mechanisms that drive PDAC metastasis and identify actionable therapeutic vulnerabilities.

Activating mutations in KRAS occur in approximately 90% of PDAC cases and represent the principal oncogenic driver of PDAC ^12^. The most prevalent variant, KRAS^G12D^, which confers the worst oncologic outcomes results in constitutive activation of downstream effectors such as the MAPK/ERK and PI3K/AKT signaling pathway, leading to autonomous tumor cell proliferation and survival ^13–16,42^. KRAS activation also promotes invasion and metastatic dissemination through cytoskeletal remodeling and enhanced migratory capacity. While recent therapeutic advances such as small-molecule KRAS inhibitors have generated optimism ^7–11^, resistance inevitably develops, highlighting the urgent need for complementary or alternative targets that modulate KRAS-driven signaling and metastatic tumor behavior.

The actin-regulatory protein Mena (ENAH), a member of the Ena/VASP family, has emerged as a critical mediator of cancer cell motility and dissemination resulting in metastasis across several tumor types ^17–26^. Mena family splicing variants regulate actin filament elongation through their proline-rich EVH1, and EVH2 domains, and inhibition of phosphatases to regulate downstream kinase pathway signaling through their MenaINV and Mena11a exons facilitating cellular protrusion, migration, and invadopodia formation ^26–33, 48^. Tumor cells exploit these functions to enhance dissemination and metastatic colonization, and high Mena expression correlates with poor outcomes in gastric, hepatic, oral, and breast cancers ^19,20,24,34,35^. These findings suggest that Mena promotes the oncogenic functions of multiple driver mutations. However, its role in KRAS-mutant PDAC, particularly in the context of KRAS^G12D^-driven disease, remains largely unexplored.

Here, we demonstrate that Mena expression is heterogeneous in human and murine PDAC, correlates with reduced patient survival, and enhances migration, invasion, proliferation, and metastasis in KRAS^G12D^-mutant PDAC through modulation of ERK and AKT phosphorylation *via* phosphatase regulation. These findings identify Mena as a potential therapeutic target and mediator of KRAS-driven tumor cell signaling in PDAC.

## MATERIALS AND METHODS

### TCGA analysis

PanMena (ENAH) mRNA expression and clinical data were obtained from TCGA PDAC cohorts. Overall survival was analyzed using Kaplan–Meier methods with log-rank testing. Patients were stratified by median ENAH expression.

### Cell lines

A validated murine PDAC KPC (KRAS^G12D+/-^/Tp53^R172H+/-^/Pdx-1-Cre) cell line was purchased from Cancer Research UK Glasgow: The Beatson Institute (Ximbio/CancerTools catalog #153474, Jennifer Morton Laboratory) ^36^. Panc02 (Smad4 *df*) cell line was purchased commercially (NCI-DTP Cat# PAN 02). Cells were cultured under standard conditions. Validated short-hairpin RNA (Sigma Aldrich #SHCLND, MISSION shRNA Clone ID TRCN0000312163) against the panMena transcript (Gene ID 13800, CDS) was directly transfected to generate shMena cell lines. Selection for puromycin resistance was used (10ug/mL).

A Mena Classic overexpression lentiviral expression vector was generated in-house by modifying a pLenti CMV GFP Hygro (addgene Plasmid #17446, RRID: Addgene_17446) backbone by removing GFP and inserting mouse Mena cDNA (CDS: 66…1691, Genebank #BC062927). Selection for hygromycin resistance was used (1mg/mL). Sequences and constructs outlined in Figure S2.

KPC and Panc02 cells modified for Mena expression were transfected to express firefly Luciferase plasmid (Luciferase-pcDNA3, addgene Plasmid #18964, William Kaelin Lab). Antibiotic selection was performed using G418 (1mg/mL).

### Western blotting

Cells at 70-80% confluency were PBS washed and lysed on ice with protease/phosphatase inhibitor cocktail per reagent protocol (Cell Lysis Buffer II, FNN002; Halt Protease and Phosphatase Inhibitor Cocktail 100X, 78440). Protein concentration was quantified (Pierce, #23225) and 20ug loaded per well (NuPAGE Bis-Tris Mini Protein Gels, NP0321BOX). Gels were run at 100V (NuPAGE MES SDS Running Buffer 20X, NP0002) and transferred to nitrocellulose membrane via 20 minute, 20V semi-dry transfer (10X Tris/CAPS Buffer, Bio-Rad #1610778). Membranes were washed in TBS-T and blocked for 1 hour using 3% BSA in TBS-T. Primary antibodies were diluted in 3% BSA and incubated overnight at 4 degrees. Signal from LI-COR fluorescent secondary antibodies was measured using the LI-COR Odyssey CLx. Signal intensity was quantified using ImageJ and each protein normalized to GAPDH (GAPDH Loading Control GA1R, MA5-15738). Following staining for phosphorylated proteins, blots were stripped (ReBlot Plus Strong Antibody Stripping Solution, 10X, Millipore 2504) and reprobed for total protein following the above protocol. The following antibodies and dilutions were utilized in this study: Anti-Mena (panMena antibody binds all Mena isoforms), clone A351F7D9, Sigma Aldrich MAB2635; Anti-Raf-1, R&D Systems MAB4540; Anti-Phospho-Raf-1, R&D Systems PPS060; Anti-MEK1/2, Thermo-Fisher Scientific MA5-38123; Anti-Phospho-MEK1/2, Thermo Fisher Scientific PA55-117184; Anti-ERK1/2, Thermo Fisher Scientific 13-6200; Anti-Phospho-ERK1/2, Thermo Fisher Scientific 44-680G; Anti-RAS G12D Mutant Specific, Cell Signaling #14429; Anti-Phospho-AKT1, Invitrogen 44-621G; Anti-AKT1+2+3, abcam 179463; Anti-SHIP2, Cell Signaling #2839.

### *In vitro* migration and invasion

Porous (0.8uM) transwell was placed into an FBS serum gradient of 0.1-10% FBS. 50,000 KPC cells were plated for 8 hours. To observe tumor cell invasion, a layer of Matrigel was added. In both cases at the set timepoint, transwell undersides were DPBS washed and fixed with crystal violet reagent + 4% paraformaldehyde. 10, 10X fields per transwell were taken on the Leica Thunder Digital light microscope and analyzed using a custom ImageJ macro.

### *In vitro* proliferation and viability

5,000 KPC cells were plated in 24-well plates in 10% FBS DMEM and counted using CellDrop Brightfield (DeNovix Inc.) at 24-hour intervals. To quantify cell viability, Trypan blue exclusion was utilized. 24, 48, and 72 hours post-seeding of cells, medium (containing unadhered cells) and adhered cells were collected. Trypan blue was added at a 1:1 ratio and viability was measured using CellDrop Brightfield.

### Matrix degradation assay

Matrix degradation assay was performed as previously published ^37^. In summary, fluorescently labeled (568) fibronectin is plated on gelatin and glutaraldehyde to create a matrix. KPC cells with varying MENA expression levels were plated on the matrix for 72 hours. Matrices are imaged for fluorescence using the ZeissObserver CLEM microscope and 30 high-powered fields are captured at 63X. Degradation holes were quantified using a custom macro on ImageJ.

### siRNA

ON-TARGETplus SMARTpool siRNA targeting Inppl1 (SHIP2) was purchased from Dharmacon (L-040673-00-0005) with respective Non-targeting Pool (D-001810-10-05). Following standard protocol, KPC cells were electroporated with siRNA using Cell Line Nucleofector Kit V (LONZA Cat. #VCA-1003) following kit protocol. Cells were incubated with siRNA for 72 hours following optimization and validation by western blotting for SHIP2.

### XTT cell proliferation assay

To measure cell proliferation, XTT assay was used (Cayman Chemical, Item No. 10010200). Per manufacturer’s protocol, 1×10^3^ cells were plated per well in a 96-well plate, 24 hours post-nucleofection. Reagents were added 48 hours post-plating, incubated for 2 hours, and read at 450nm.

### Microscopy and image analysis

LI-COR Odyssey CLx, Leica Thunder Digital light microscope, Zeiss AxioObserver CLEM were used as described. All images were processed and analyzed using ImageJ. Custom macros were generated to quantify cell invasion, migration, and degradation spots. Protocols for imaging were developed with the support of the Analytical Imaging Facility.

### *In vivo* orthotopic model

6,000 cells per injection were trypsinized and resuspended in 50uL of PBS. Following a previously published aseptic procedure, mice pancreata were delivered into a left subcostal wound and directly injected with a single cell line per syngeneic C57/B6J mouse (Jackson Labs) at 8-10 weeks of age ^38,39^. As PDAC afflicts both females and males, both genders of mice were randomized into groups and used. Pancreata were placed back *in situ* into the abdominal cavity and the abdominal wall and skin were sutured closed. Splenic injection model followed previously published protocol, utilizing 50,000 cells per injection, and the described surgical technique ^40^. All animal models utilized in this study were IACUC approved under Albert Einstein College of Medicine (AECOM) Protocol #00001079 and #00001845.

### Tumor Volume

During tumor resection, adjacent normal pancreas and spleen are cleared. Length and width of each tumor is measured using digital caliper. Tumor volume was measured as (length)x(width)^2^/2.

### *In Vivo* Imaging System (IVIS)

Luciferase-expressing cells were validated for bioluminescence upon tumor growth by orthotopic cell injection followed by Luciferin (VivoGlo^TM^ Luciferin, In Vivo Grade, Promega P1042) administration at 150mg/kg dose using the In Vivo Imaging System Facility (IVIS) at AECOM. Mice were imaged weekly following orthotopic injection for bioluminescence of tumors. Bioluminescence was quantified per tumor using Living Image software provided by AECOM.

### Transgenic Mouse Model

The KPC transgenic mouse strain (KRAS^G12D(LSL–KRAS)^; Trp53^R172H (LSL-Trp53)^; Pdx1-Cre) was generously gifted by the Laboratory of Ben Stanger, University of Pennsylvania. Mice were sacrificed and pancreata were collected at 10-14 weeks (pancreatic intraepithelial neoplasia) and 16+ weeks (pancreatic ductal adenocarcinoma), formalin fixed, paraffin-embedded (as described below) and stained for panMena expression.

### Human specimens

PDAC samples from patients who underwent curative-intent surgical resection between 2013 and 2022 at a single tertiary care institution were included for this study (n=60, IRB-2018-8906). These patients included those who received neoadjuvant chemotherapy as well as those that were treatment-naïve at the time of surgery. For each patient, the diagnosis of PDAC was confirmed.

### Tissue processing and pathological analysis

Murine tissue was processed by the AECOM Histology and Pathology Core Facility. All tissue was harvested and immediately fixed in 10% formalin for 24 hours, followed by paraffin embedding. Formalin-fixed paraffin-embedded (FFPE) tissue was mounted to glass slides in 2uM sections. Slides were stained by hematoxylin & eosin (H&E) to confirm tumor presence. All pathological analyses were performed by a board-certified gastrointestinal pathologist (P.P.) using the Nikon Eclipse Ci microscope and Adobe Photoshop 2025 for image analysis.

### Immunohistochemistry

Sectioned tissue from murine PDAC primary and secondary tumors harvested from orthotopically injected or transgenic mice were stained using immunohistochemistry for panMena. Standard immunohistochemical protocol was followed using ImmPRESS HRP Horse Anti-Mouse kit (MP-7402). In brief, FFPE slides were deparaffinized and antigen retrieval was performed using pH 6 citrate buffer. Tissue was blocked using BLOXALL Blocking solution followed by 2.5% normal horse serum. Primary antibody (Anti-Mena, 1:100) was applied overnight at 4 degrees. Tissue was incubated with ImmPRESS Polymer reagent followed by peroxidase substrate solution. Slides were then hematoxylin counterstained, cleared, and mounted. Orthotopic tumors were stained for Ki-67 (Anti-Ki-67, D3B5; #12202) following the same outlined protocol. All immunohistochemistry was scored by board-certified pathologist (P.P.). Patient samples were stained using a validated immunohistochemical staining protocol for TMEM doorways, as previously described ^19,20^.

### Statistical analysis

GraphPad Prism was utilized to perform statistical analyses. Student’s t-tests were utilized in all cases to compare overexpression or knockdown cells to their respective controls. In the case of comparing multiple groups, Two-Way ANOVA was used. In the case of correlating survival outcomes, Log-rank Kaplan-Meier analysis was used.

### Data availability

Mena expression survival data were obtained from The Cancer Genome Atlas data extracted from the Protein Atlas (Expression of ENAH in pancreatic cancer – The Human Protein Atlas). All other data and detailed protocols are available upon request from the corresponding author.

### Ethics statement

All animal studies were approved by the Institutional Animal Care and Use Committee (IACUC) at Albert Einstein College of Medicine (Protocol #00001079 and #00001845). Human tissue studies were approved by the Institutional Review Board (IRB-2018-8906). All quantitative analysis was performed blinded to experimental groups and/or using semi-automated software.

## RESULTS

### High Mena expression correlates with poor survival in patients with locoregional PDAC

Given the established role of Mena in tumor invasion and metastasis across multiple cancer types¹ ^-^², we evaluated whether Mena expression correlates with outcomes in PDAC. Analysis of TCGA data from 176 patients with PDAC — predominantly locoregional disease (Fig. 1A) — demonstrated that high panMena mRNA expression was associated with significantly worse overall survival compared to low-expression tumors (median OS 1.5 vs 3.1 years; p = 0.0047; Fig. 1B). As most patients with locoregional PDAC ultimately succumb to metastatic progression, these findings suggest a functional link between Mena expression and metastatic lethality.

**Figure 1.**
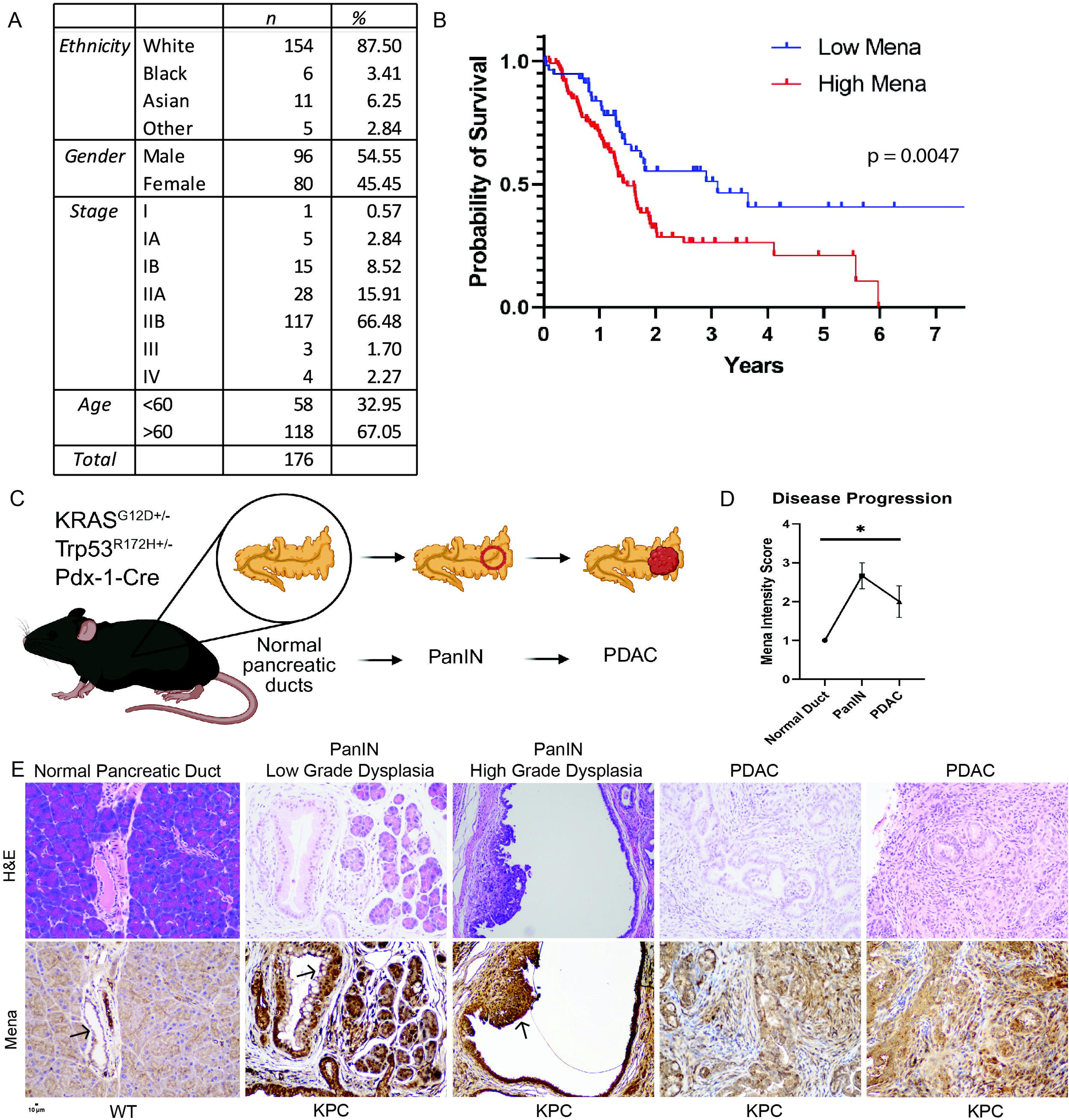
Mena expression promotes tumor initiation and progression. **A.** Patient demographics and staging information. Patients were primarily diagnosed with locoregional disease. **B.** Patients were grouped as high or low panMena-expressing and survival plotted. Low Mena expression (n=59), High Mena expression (n=117); p = 0.004, Log-rank (Mantel-Cox). **C.** KPC (KRAS^G12D+/-^/Tp53^R172H+/-^/Pdx-1-Cre) transgenic mice progression through clinical stages of PDAC development. **D.** Mena intensity score as scored by pathologist (P.P) in ductal cells over PDAC progression, p=0.03; two-way ANOVA. **E.** H&E stained (top row) and panMena stained (bottom row) pancreas specimen from left to right: normal pancreatic duct (WT pancreas), PanIN low grade dysplasia (KPC transgenic mouse), PanIN high grade dysplasia (KPC transgenic mouse), PDAC (KPC transgenic mouse); identified by board-certified pathologist (P.P.). Arrows indicate pancreatic ductal cells in normal and PanIN-afflicted pancreata. panMena expression intensity is increased in pre-invasive PanIN lesions in a KPC transgenic mouse model compared to normal ductal cells and PDAC, n=3.

To assess Mena protein expression in human tumors, we analyzed an institutional PDAC cohort (n = 60; Fig. S1). Mena protein was heterogeneously expressed both between and within tumors, however, all samples exhibited detectable Mena expression (Fig. S1A). Mena expression did not differ significantly between treatment-naïve tumors and those exposed to neoadjuvant chemotherapy (Fig. S1C), indicating that Mena expression is maintained despite cytotoxic treatment.

### Mena promotes tumor initiation and PDAC progression in KRAS^G12D^ transgenic mice

To determine whether Mena expression changes during PDAC development, pancreata from KPC (KRAS^G12D+/−^; Trp53^R172H+/−^; Pdx1-Cre) mice were examined across disease stages (Fig. 1C) ^36,41^. Normal pancreatic ducts exhibited low Mena expression, whereas both low- and high-grade PanIN lesions displayed significantly increased Mena staining intensity (Fig. 1D–E, p=0.03). Elevated Mena expression was maintained in invasive PDAC lesions, indicating that Mena upregulation is an early and sustained feature of KRASG12D-driven tumorigenesis.

### Mena modulation in KRAS^G12D^ PDAC model regulates migration, invasion, and matrix degradation *in vitro*

Previous reports implicate that co-option of Mena signaling by tumor cells promotes migration and invasion processes, resulting in increased tumor cell dissemination ^18,20,26^. To assess the functional role of Mena in KRAS-mutant PDAC, murine KPC cells were engineered to stably overexpress Mena Classic (MenaOE) or undergo panMena knockdown (shMena), with vector controls (Fig. S2). Mena modulation was confirmed by western blotting (Fig. 2A). Mena Classic is the most abundant isoform of Mena and therefore is the focus of the study. In transwell assays, MenaOE cells exhibited significantly increased migration and invasion, whereas Mena knockdown markedly reduced both processes (Fig. 2B–D). Consistent with enhanced invasive capacity, MenaOE cells demonstrated increased matrix degradation, while shMena cells showed reduced degradative activity (Fig. S3B). These findings indicate that Mena Classic amplifies KRAS^G12D^-driven invasive behavior in PDAC cells.

**Figure 2.**
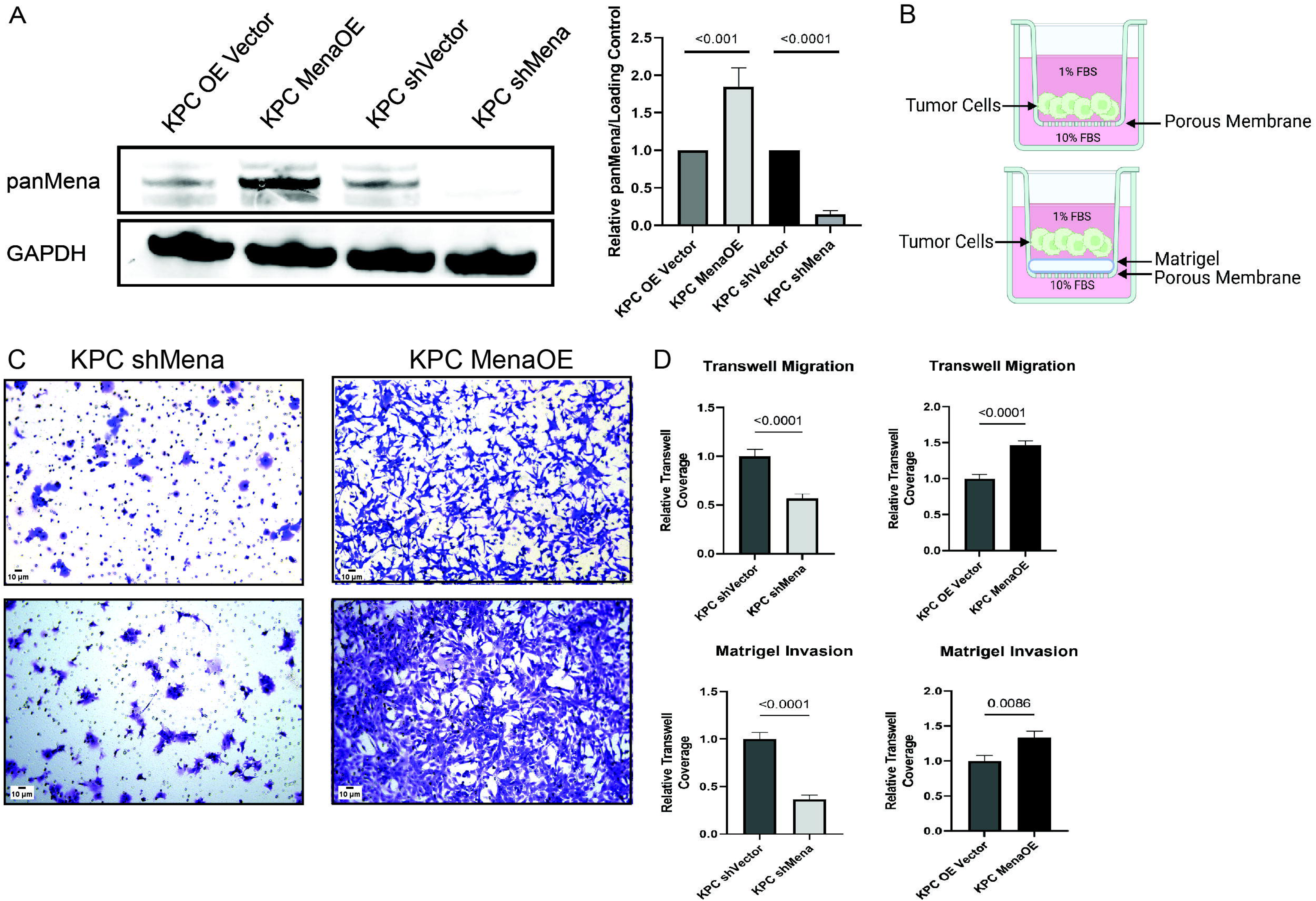
Mena promotes KRAS-mediated migration and invasion. **A.** A stable cell line derived from a KRAS^G12D+/-^/Tp53^R172H+/-^/Pdx-1-Cre transgenic mouse was utilized as the model of interest. Modified cell lines were validated for knockdown and overexpression of Mena protein by western blotting using panMena antibody (left panel). Quantified panMena intensity by western blotting (right panel), KPC shMena vs KPC shVector p=<0.0001, KPC Mena Overexpression vs KPC Overexpression Vector p=<0.001. **B.** Schematic image of transwell migration (upper panel) and transwell invasion (lower panel) experimental set-up prior to transmigration or invasion of tumor cells. **C.** Representative images of transwell underside following 8 hours of tumor cell migration of KPC shMena (top, left panel), KPC Mena Overexpression (top, right panel) and tumor cell invasion overnight of KPC shMena (bottom, left panel), KPC Mena Overexpression (bottom, right panel). Tumor cells which have transmigrated or invaded to the underside of the transwell are stained using Crystal Violet reagent and visualized in purple. **D.** Upper panels: Plotted quantification of tumor cell coverage on migration transwell underside (top panel). 10 HPFs per transwell were processed using custom ImageJ macro; KPC shMena vs KPC shVector p=<0.0001, KPC Mena Overexpression vs KPC Overexpression Vector p=<0.0001. Lower panels: Plotted quantification of tumor cell coverage on invasion transwell underside (bottom panel). 10 HPFs per transwell were processed using custom ImageJ macro; KPC shMena vs KPC shVector p=<0.0001, KPC Mena Overexpression vs KPC Overexpression Vector p=0.0086. All statistical analyses were performed using student’s t-tests.

### Mena regulates primary tumor growth *in vivo* in KRAS^G12D^ PDAC

To determine whether Mena regulates tumor growth in vivo, modified KPC cells were orthotopically injected into the pancreas (Fig. 3A). By 14 days post-injection, MenaOE cells formed large pancreatic tumors, whereas shMena cells failed to establish palpable tumors, demonstrating delayed tumor initiation (Fig. 3B). Quantitative analysis confirmed significantly increased tumor volume in MenaOE tumors relative to controls (Fig. 3C).

**Figure 3.**
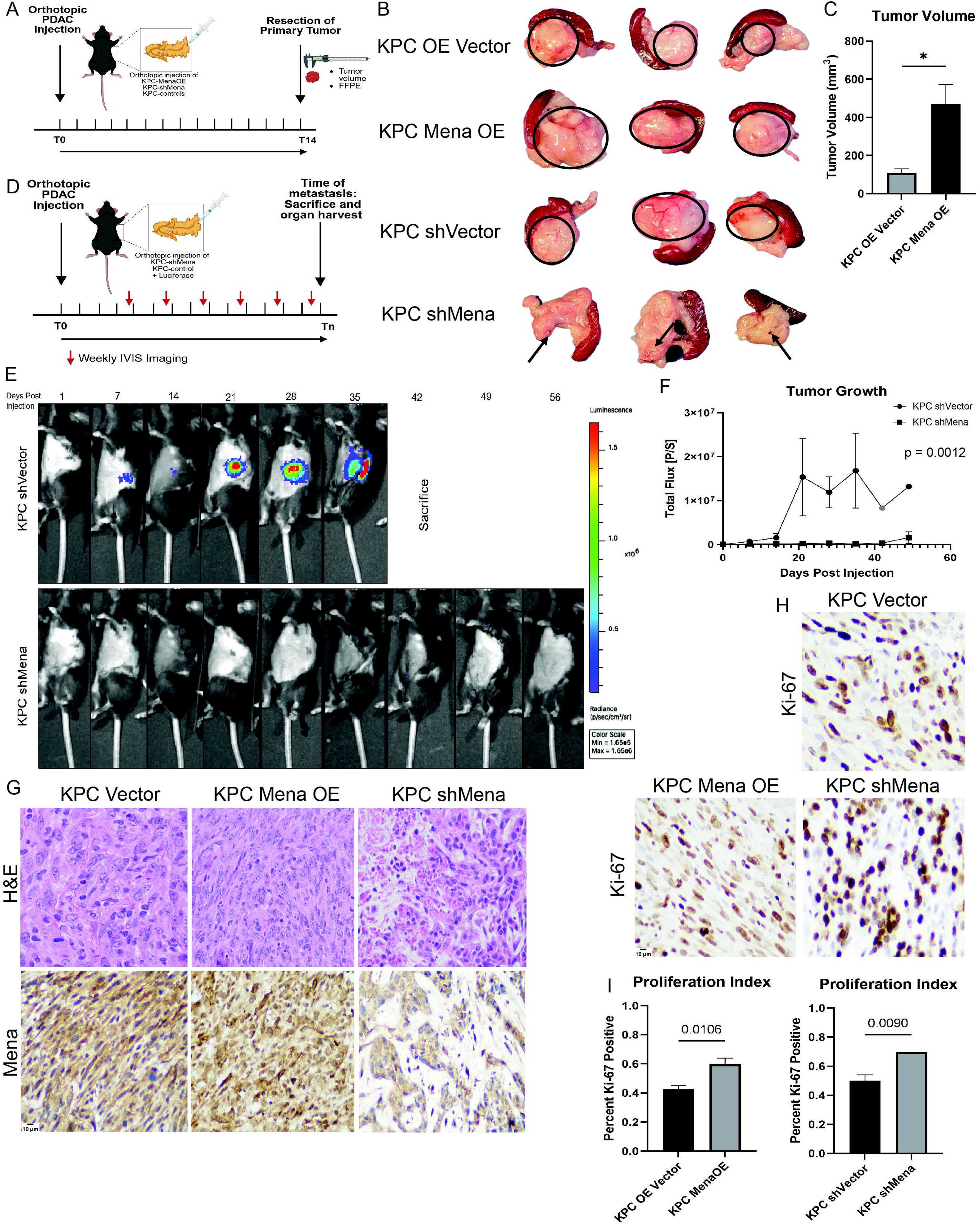
Mena regulates KRAS-driven primary tumor growth *in vivo.* **A.** Schematic of orthotopic injection model. 6,000 tumor cells are injected directly into mouse pancreata during aseptic survival surgery. 14 days post-injection, tumors are resected and analyzed by various metrics. **B.** Representative images of tumor size with adjacent normal pancreas 14 days post-injection. Tumor areas are outlined by black circles. In the case of no visible tumor, injection site is indicated by arrow. **C.** Quantified tumor volume using the following formula: (length)x(width)^2^/2; KPC OE Vector vs KPC MenaOE, p=0.0145. **D.** Schematic of orthotopic injection model with luciferase-tagged KPC shVector and shMena cell lines using In Vivo Imaging System (IVIS). **E.** Weekly tumor bioluminescence imaging measured using radiance (Total Flux p/sec). **F.** Quantified Total Flux p/sec over time. Total Flux represents luciferin uptake by luciferase expressing tumor cells and is a surrogate metric for tumor size. Gray point represents timepoint with outlier removed; KPC shVector vs KPC shMena, p = 0.0012, Mixed-effects analysis. **G.** H&E and IHC slide of primary tumor areas stained for Mena expression to determine that Mena expression is maintained in tumor cells following in vivo tumor growth; KPC Vector (left panel), KPC Mena Overexpression (center panel), KPC shMena (right panel). **H.** Representative Ki-67 staining of tumor areas; KPC Vector (left panel), KPC Mena Overexpression (center panel), KPC shMena (right panel). **I.** Proliferation Index is defined as percent Ki-67 positivity as quantified by board-certified pathologist (P.P). KPC Mena Overexpression vs KPC Overexpression Vector p = 0.0106, KPC shMena vs KPC shVector p = 0.0090. Statistical analyses were performed using student’s t-tests except when noted otherwise.

Longitudinal bioluminescence imaging of luciferase-tagged shVector and shMena cells revealed delayed tumor detection and significantly reduced tumor burden over time in the absence of Mena (Fig. 3D–F). Tumors that eventually formed in shMena-injected mice retained panMena expression (Fig. 3G), suggesting selective outgrowth of Mena-positive cells. Differences in tumor growth were not attributable to altered cell viability (Fig. S4). Ki-67 staining demonstrated increased proliferation in tumors which overexpressed Mena Classic or recovered Mena expression (Fig. 3H–I), indicating that Mena promotes KRAS-driven tumor growth.

### Mena regulates metastatic tumor growth, metastasis frequency, and *in vivo* survival in KRAS^G12D^ PDAC-bearing mice

Following orthotopic tumor establishment, mice underwent curative-intent distal pancreatectomy and were monitored for metastatic progression. Mice bearing MenaOE primary tumors developed spontaneous liver metastases, whereas shMena tumors produced infrequent and significantly smaller metastatic lesions (Fig. 4A; Fig. S5B). Metastases derived from MenaOE tumors maintained high panMena expression (Fig. 4B).

**Figure 4.**
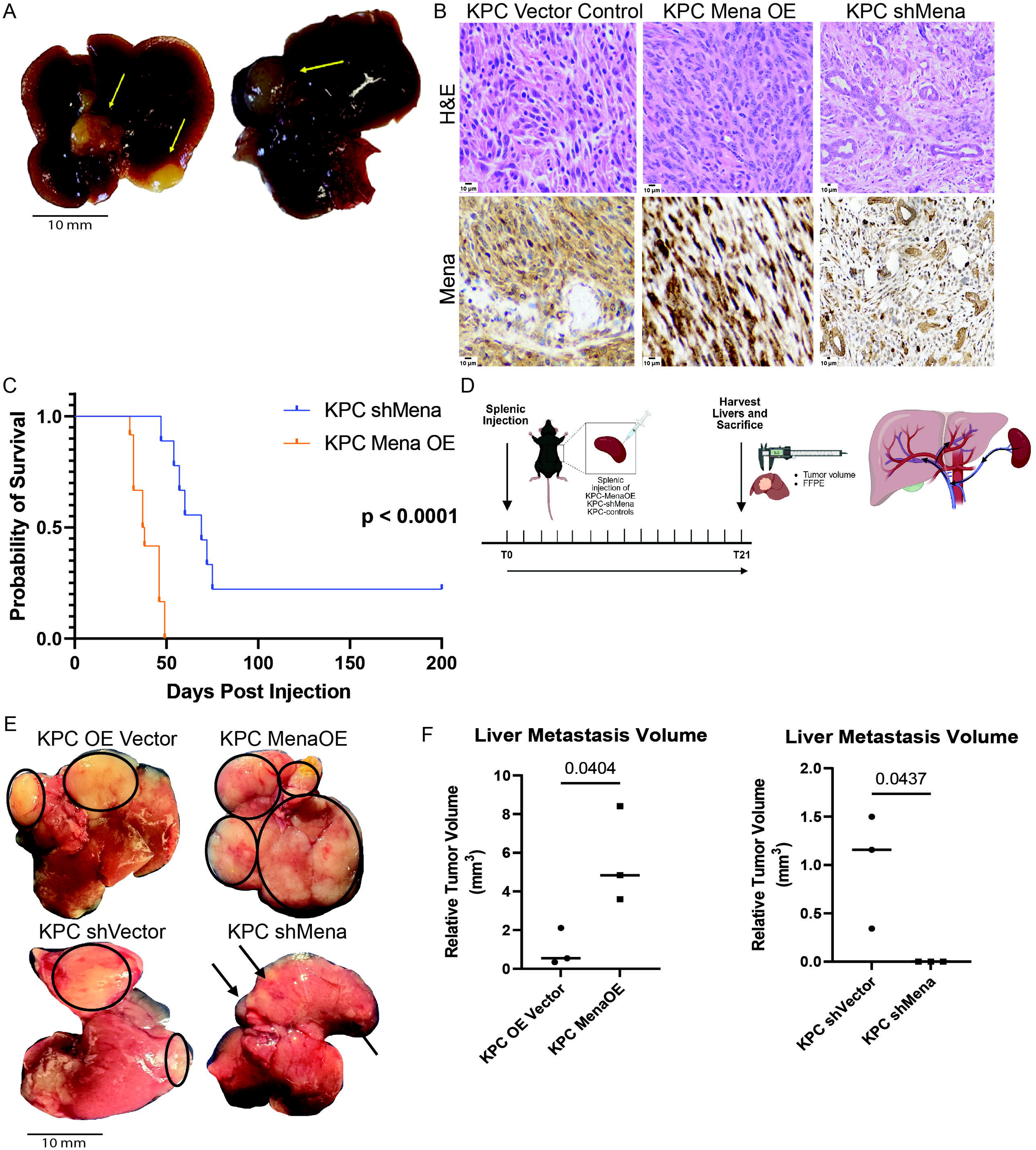
Mena regulates metastatic tumor growth, metastasis frequency, and survival *in vivo.* **A.** Representative images of spontaneous liver metastases from KPC Mena Overexpression primary tumors. Arrows indicate macrometastases visible in harvested liver specimen. **B.** H&E and IHC slides of metastatic tumor areas stained for panMena expression; KPC Vector (left panel), KPC Mena Overexpression (center panel), KPC shMena (right panel). **C.** Survival plot comparing PDAC-bearing KPC Mena Overexpression and KPC shMena injected mice, p < 0.0001; Median survival 37.5 vs 69 days post-injection. **D.** Schematic outline of experimental metastasis splenic injection model. **E.** Representative images of liver metastases resulting from splenic injection (experimental metastasis, direct seeding of liver). Metastatic foci are outlined in black or indicated by arrows. In the case of MenaOE, entire liver was colonized by macro metastases. **F.** Relative tumor volume of liver metastases. Metastatic tumor volume was measured as (length)x(width)^2^/2; KPC Mena Overexpression vs KPC Overexpression Vector, p = 0.0402; KPC shMena vs KPC shVector, p = 0.0437.

Consistent with increased metastatic burden, mice bearing MenaOE tumors exhibited significantly reduced survival compared to mice bearing shMena tumors (median survival 37.5 vs 69 days; p < 0.0001; Fig. 4C; Table S1).

As Mena isoforms are known to play a significant role in tumor cell dissemination ^20,26,32,33^, we sought to separate Mena Classic’s effect on dissemination from metastatic growth. PDAC cells were injected intrasplenically (Fig. 4D). MenaOE cells rapidly colonized the liver, producing extensive metastatic disease, whereas shMena cells formed only small metastatic foci (Fig. 4E). Quantitative analysis confirmed significantly increased metastatic tumor volume in MenaOE cells and reduced volume in shMena cells (Fig. 4F), demonstrating that Mena Classic also regulates metastatic outgrowth.

### Mena sustains ERK and AKT signaling downstream of KRAS^G12D^ and WT KRAS

Given the observed effects on tumor growth, we investigated whether Mena modulates KRAS signaling pathways. Under serum starvation, Mena knockdown significantly reduced ERK1/2 phosphorylation, with modest effects on upstream RAF and MEK activation (Fig. S6; Fig. 5A–B). AKT phosphorylation was also significantly reduced in shMena cells (Fig. 5C–D).

**Figure 5.**
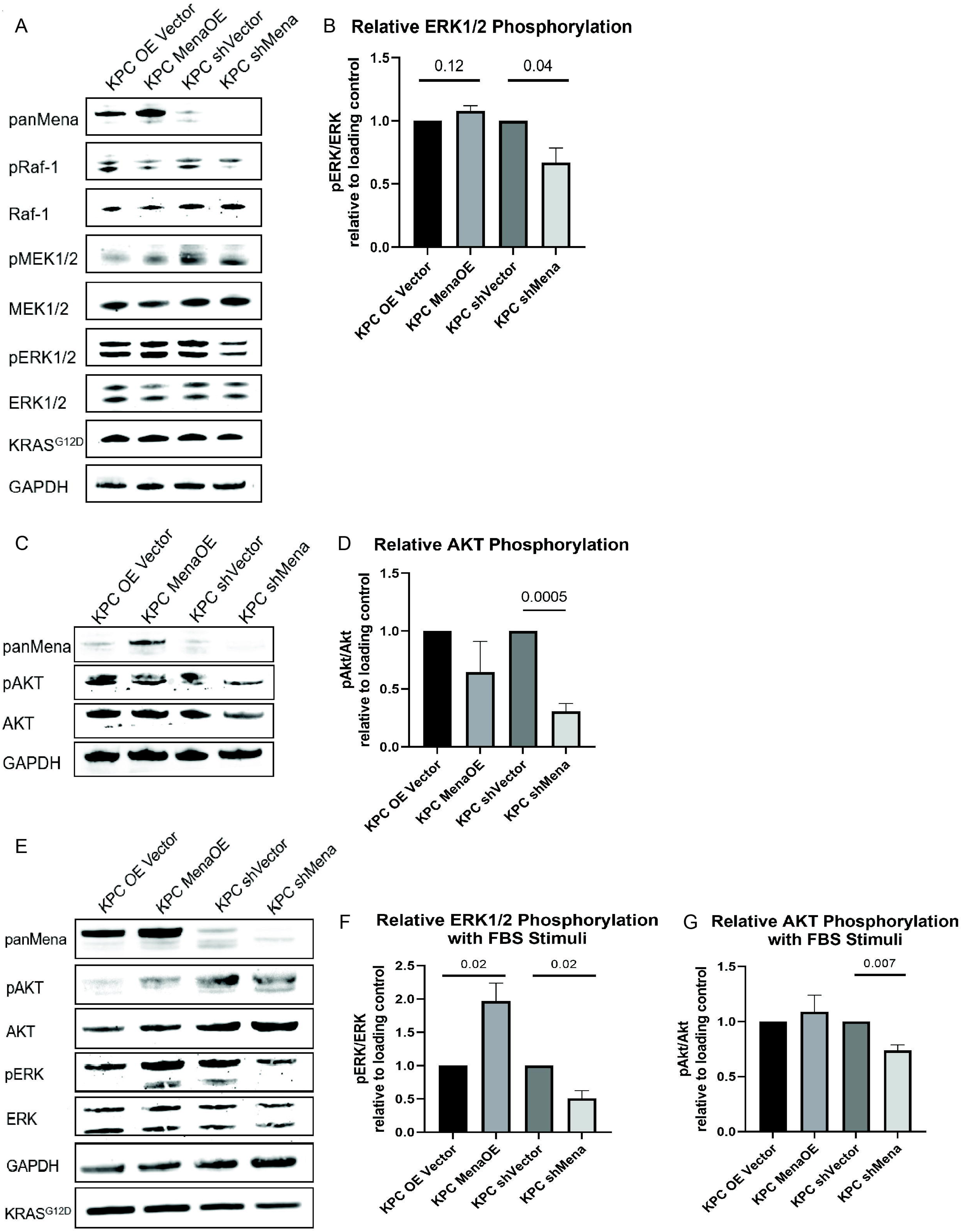
Loss of Mena decreases ERK1/2 and AKT activity downstream of KRAS^G12D^. **A.** Western blotting for panMena, Raf-1, phospho-Raf-1, MEK1/2, phosphor-MEK1/2, ERK1/2, phospho-ERK1/2, KRAS^G12D^, and GAPDH in each cell line of interest with varying levels of Mena expression, following 24 hours of starvation conditions. **B.** Phosphorylated protein was measured relative to total protein as a metric of kinase activity. Phosphorylated protein and total protein were both quantified relative to loading control. KPC Mena OE and KPC shMena relative protein levels were compared to normalized control vector cell lines, per blot, respectively. Phosphorylated ERK1/2 slightly increases in Mena overexpressing cells and is significantly decreased in Mena knockdown cells (KPC Mena Overexpression vs KPC Overexpression Vector, p=0.14; KPC shMena vs KPC shVector, p=0.04). **C.** Western blotting for panMena, phospho-AKT, AKT, and GAPDH, following 24 hours of starvation conditions. **D.** Phosphorylated protein was measured relative to total protein, after normalization to loading control. KPC Mena OE and KPC shMena relative protein levels were compared to normalized control vector cell lines, per blot, respectively. Phosphorylated AKT significantly decreases in Mena knockdown cells (KPC shMena vs KPC shVector, p=0.0005). Phosphorylated AKT is not significantly changed in MenaOE cells. **E.** Western blotting for panMena, phospho-AKT, AKT, phospho-ERK1/2, ERK1/2, GAPDH, KRAS^G12D^, in FBS stimulated conditions. **F.** Phosphorylated ERK1/2 significantly increased in Mena overexpressing cells and significantly decreased in Mena knockdown cells (KPC Mena Overexpression vs KPC Overexpression Vector, p=0.02; KPC shMena vs KPC shVector, p=0.02). **G.** Phosphorylated AKT significantly decreased in Mena knockdown cells (KPC shMena vs KPC shVector, p=0.007).

Upon receptor tyrosine kinase stimulation, ERK1/2 phosphorylation increased in MenaOE cells but remained suppressed in shMena cells (Fig. 5E–F). AKT phosphorylation similarly remained dependent on Mena expression (Fig. 5G). These findings indicate that Mena sustains ERK and AKT activity downstream of both mutant and wild-type KRAS. These findings are supported by previous results which show KRAS^G12X^ cells signal along the PI3K/AKT axis through their mutant KRAS allele which is enhanced by RTK stimulation. ^45^

To assess pathway specificity, Mena expression was modulated in Panc02 (SMAD4-deficient) PDAC cells. In this non-KRAS-driven context, Mena overexpression or knockdown did not alter ERK or AKT phosphorylation (Fig. 6A–C) or tumor growth in vivo (Fig. 6D–F), demonstrating that Mena functions as a KRAS-dependent signaling regulator.

**Figure 6.**
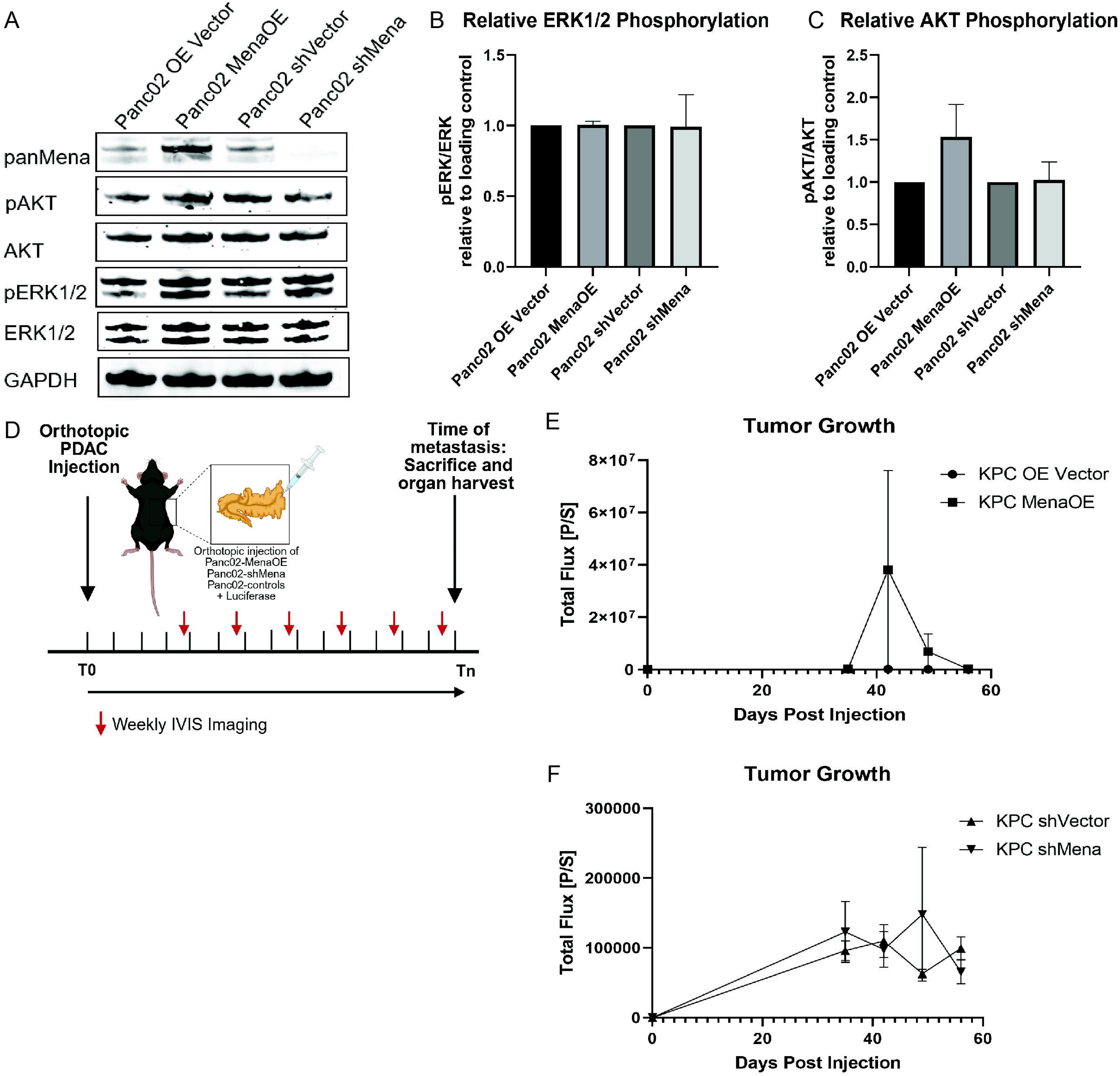
Smad4 mutant PDAC cells’ ERK1/2 and AKT activity are not regulated by Mena. **A.** Western blotting for panMena, phospho-AKT, AKT, phospho-ERK1/2, ERK1/2, GAPDH in Panc02 (Smad4 *df*) cells modified for Mena expression following 24 hours of starvation. **B.** ERK1/2 phosphorylation is unchanged by Mena overexpression or Mena knockdown in Panc02 cells. **C.** AKT phosphorylation is unchanged by Mena overexpression or Mena knockdown in Panc02 cells. **D.** Schematic of orthotopic injection model with luciferase-tagged Panc02 Vector controls, MenaOE, shMena cell lines using In Vivo Imaging System (IVIS). **E.** In a non-KRAS mutant background, Mena does not regulate tumor growth as demonstrated by Quantified Total Flux p/sec over time. Panc02 OE Vector vs Panc02 MenaOE, p = 0.3141, Two-way ANOVA. **F.** Quantified total flux over time. Panc02 shVector vs Panc02 shMena, p = 0.5661, Two-way ANOVA.

### Mena promotes ERK and AKT activity via SHIP2 regulation

To determine the signaling mechanism underpinning Mena’s contribution to tumor cell proliferation, we hypothesized that Mena regulates phosphatases to modulate ERK1/2 and AKT activity. SHIP2 is one known phosphatase which is a binding partner of Mena ^27,47^. To determine whether SHIP2 mediates Mena-dependent signaling, SHIP2 was transiently knocked down in Mena-modified KPC cells. SHIP2 depletion in MenaOE cells further increased ERK1/2 phosphorylation and modestly increased AKT phosphorylation (Fig. 7A–C), accompanied by increased proliferative capacity (Fig.

**Figure 7.**
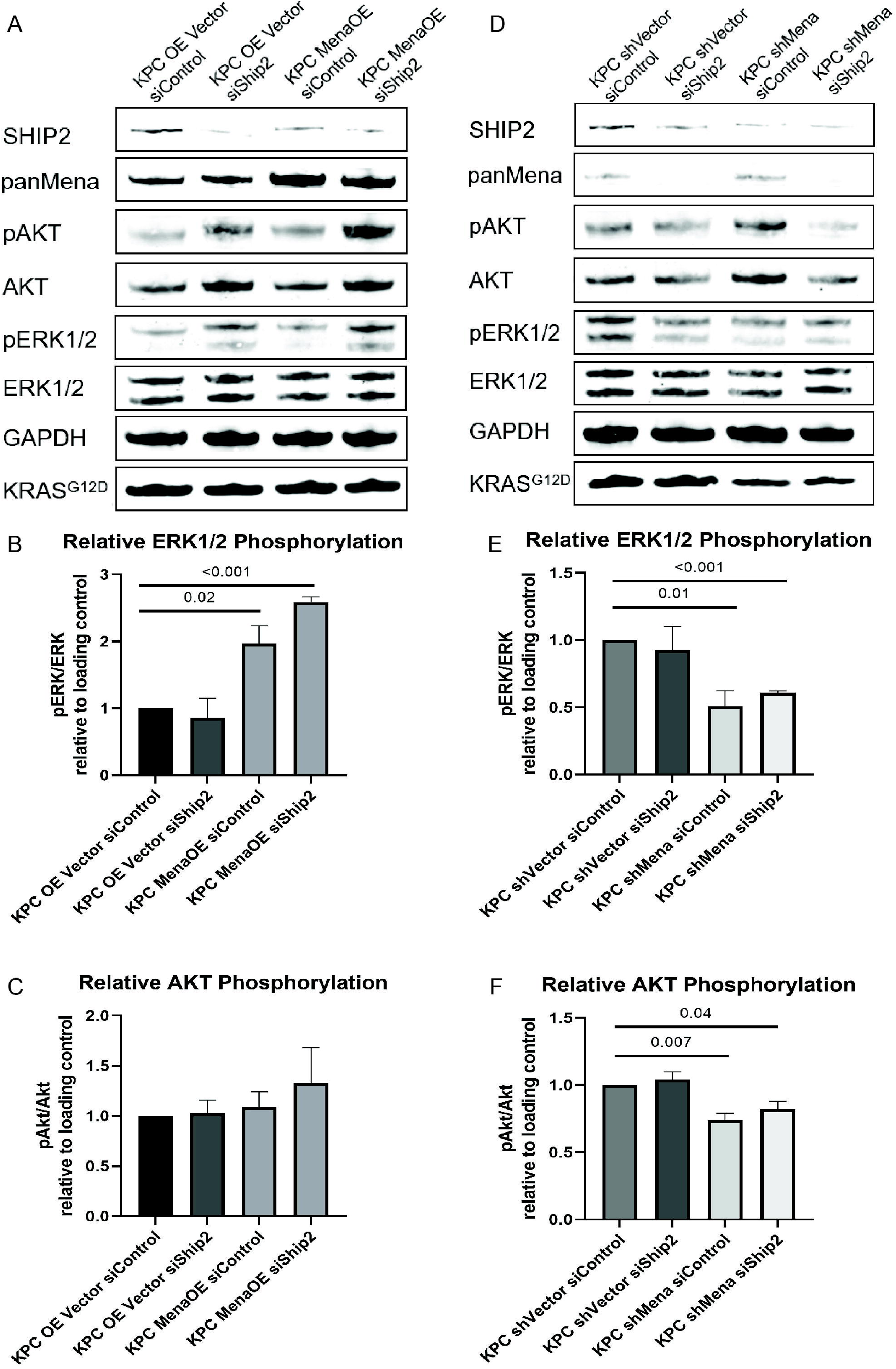
Mena acts on SHIP2 to regulate kinase activity. **A.** Western blotting for SHIP2, panMena, phospho-AKT, AKT, phospho-ERK1/2, ERK1/2, GAPDH, KRAS^G12D^. KPC Overexpression Vector and KPC Mena Overexpression cell lines were electroporated with siRNA against SHIP2 (or non-targeting). **B.** When SHIP2 is knocked down, ERK1/2 phosphorylation increases further in KPC Mena Overexpression cells compared to controls (p=<0.001). **C.** AKT phosphorylation modestly increases in KPC Mena Overexpression cells with SHIP2 knockdown. **D.** Western blotting for SHIP2, panMena, phospho-AKT, AKT, phospho-ERK1/2, ERK1/2, GAPDH, KRASG12D. KPC shVector and KPC shMena cell lines were electroporated with siRNA against SHIP2 (or non-targeting). **E.** When SHIP2 is knocked down, ERK1/2 phosphorylation remains significantly decreased in KPC shMena cells compared to controls (p=<0.001), but levels of relative phosphorylated ERK1/2 slightly increase**. F.** When SHIP2 is knocked down, AKT phosphorylation remains significantly decreased in KPC shMena cells compared to controls, but levels of relative phosphorylated AKT are nearly recovered to those of controls (p=0.04).

S7A). In shMena cells, SHIP2 knockdown only partially restored kinase phosphorylation (Fig. 7D–F), suggesting that Mena regulates ERK and AKT signaling through SHIP2 and potentially additional phosphatases. A schematic model is presented in Fig. 8.

**Figure 8.**
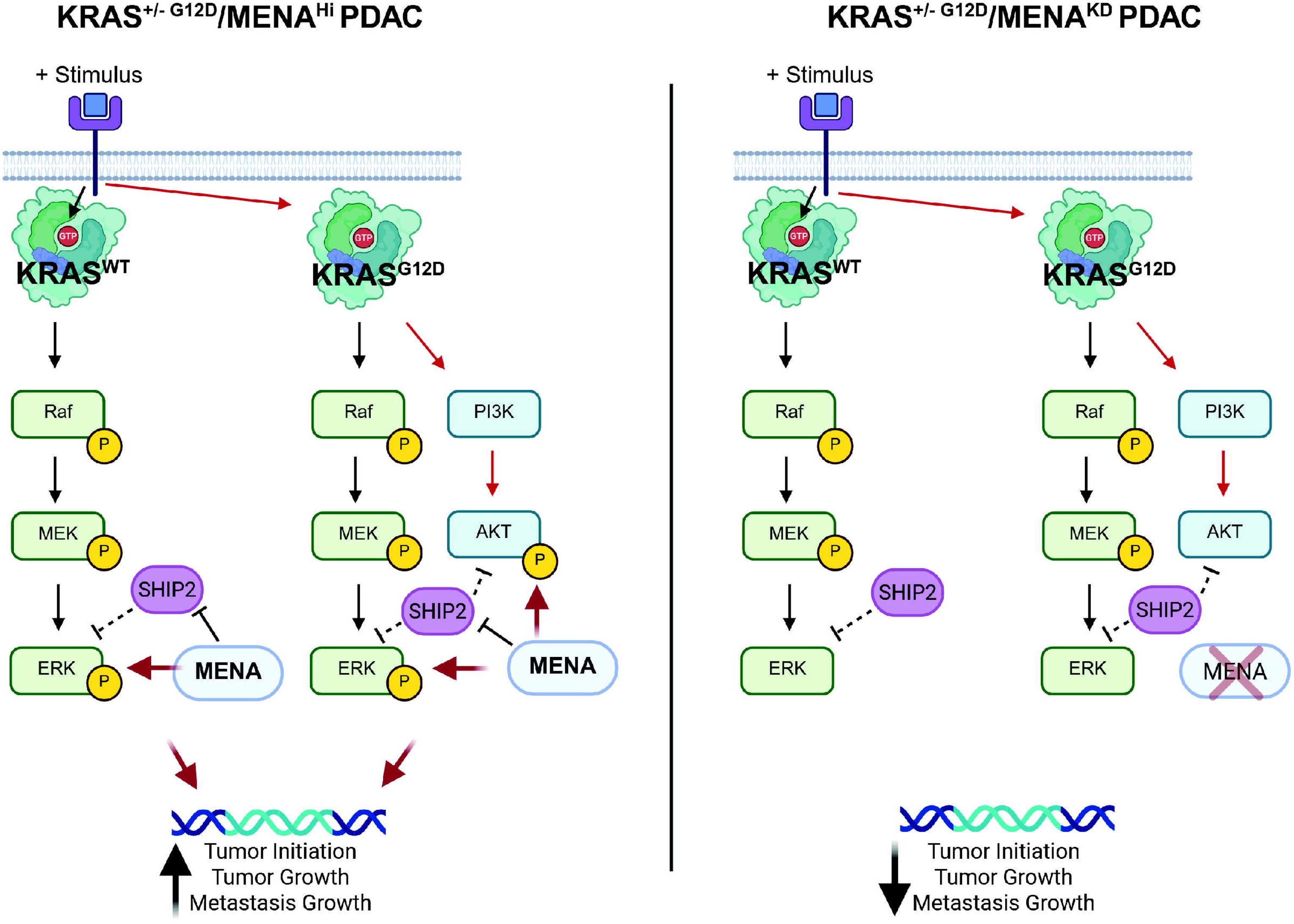
Proposed mechanism of action for Mena in modulating KRAS^G12D^ and WT KRAS signaling. Left panel: In KRAS^G12D+/-^ Mena^High^ tumor cells, Mena promotes ERK1/2 phosphorylation downstream of KRAS^G12D^ and WT KRAS through a partially SHIP2-mediated mechanism. AKT activity is maintained downstream of KRAS^G12D^ when Mena is overexpressed. Right panel: In KRAS^G12D+/-^ Mena^KD^ tumor cells, loss of Mena depletes ERK1/2 activity downstream of KRAS^G12D^ and WT KRAS, as well as AKT activity downstream of KRAS^G12D^. The effect of Mena knockdown is slightly rescued by SHIP2 knockdown, supporting that this phosphatase plays a role in Mena’s mechanism of action.

## DISCUSSION

### High levels of Mena correlate with poor outcomes in PDAC

High levels of panMena mRNA expression correlate with poor survival, supporting that Mena family proteins play an important role in metastasis, as most patients with PDAC succumb to metastasis. Survival outcomes in our mouse model reflect the patient cohort data as mice with Mena^Low^ PDAC survive approximately double the time of those with Mena^High^ PDAC. This study aimed to untangle the mechanisms behind this finding in the context of KRAS^G12D^ mutant PDAC, the primary and most aggressive subtype of PDAC^42^. We have identified that high levels of Mena Classic expression increase tumor cell migration and invasion as well as primary tumor and metastatic growth. panMena knockdown has the opposite effect, decreasing tumor cell migration, invasion, primary tumor growth, and metastasis growth. These findings show the mechanisms behind the observed survival differences in patients and mice and identify Mena as a potential prognostic factor in PDAC.

### Mena is critical in promoting KRAS^G12D^-mediated metastasis

KRAS^G12D^ PDAC which overexpresses Mena Classic results in significantly larger and more frequent spontaneous liver metastases than controls. Demonstrating that this phenotype is due to increased metastatic growth when Mena Classic is overexpressed, Mena overexpressing cells seeded directly to the liver *via* splenic injection also grow significantly larger metastases. These findings support a novel role for Mena in promoting not only primary tumor growth but also metastatic tumor growth as a mechanism to promote PDAC metastasis. Mechanistically, we show that this occurs due to ERK1/2 activity which increases when Mena is overexpressed.

The role of Mena in tumor cell migration and invasion is well-studied in breast cancer ^17,18,26,48^. As expected, we found that Mena Classic overexpression in PDAC also promotes increased migration and invasion, and panMena knockdown decreases these processes. Alternative splicing of Mena is crucial to its invasive functions. It is known that specific isoforms of Mena, such as MenaINV (+++), support invasive behavior of tumor cells and are transiently expressed, while other isoforms, such as Mena11a, support epithelial-like behavior ^18,23,27,48^. MenaINV is particularly important at TMEM doorways, which are portals for tumor cell dissemination ^18–20^. In this study, Mena Classic, the most prevalent isoform of Mena, is overexpressed, which is the classical and most abundant form of Mena. Importantly, MenaINV can be stabilized by formation of tetramers with more stable and abundant isoforms such as Mena Classic ^26^. We suspect that, combined with Mena promoting KRAS^G12D^-mediated invasion, the increased invasion and migration observed in our KPC Mena overexpression cells likely occurs through stabilization of endogenous MenaINV by the constitutively overexpressed Mena Classic. Together, these functions of Mena drive the increased metastatic burden observed in mice with Mena overexpressing, KRAS^G12D^ PDAC.

### New model of spontaneous PDAC metastasis utilizing KRAS^G12D+/-^/Mena^HIGH^ orthotopic injection

A significant challenge to the field in identifying novel targets for PDAC has been the establishment of a consistent model of spontaneous metastasis. Most orthotopic PDAC models produce gross metastases inconsistently or after long periods of time.

Transgenic KPC mouse models, while more consistent in producing spontaneous metastases, are difficult to breed and take 16-18 weeks to develop PDAC ^36,41^. Experimental metastasis models which directly seed tumor cells to sites of metastasis are often utilized to circumvent these technical challenges, however these models do not include a physiological dissemination step which is critical in tumor cell programming and therefore crucial for a biologically relevant model ^43^. When injected orthotopically, our novel KPC MenaOE cell line produced spontaneous liver metastases in 100% of injected mice ∼3.5 weeks post injection, demonstrating its consistency and biological relevance in modeling PDAC metastasis.

### Novel signaling role for Mena as a KRAS effector *via* MAPK and AKT signaling

We have identified a novel signaling role for Mena in regulating signaling downstream of KRAS^G12D^ and KRAS. Mena is necessary to maintain ERK1/2 and AKT activity downstream of both mutant and wild-type KRAS, as phosphorylation of these kinases is significantly depleted when Mena is knocked down. The presence of KRAS^G12D^ primes Mena for its regulatory function, as this effect of Mena is not seen when MAPK and AKT signaling are shunted towards other pathways. High levels of Mena in the cell promote KRAS-mediated signaling, with a particularly pronounced effect on ERK1/2; this accounts for the dramatic increase in growth of primary and metastatic tumors in the KRAS^G12D+/-^/Mena Classic^HIGH^ model.

The presence or absence of Mena in KRAS^G12D^ cells affects kinase activity through regulation of phosphatases including SHIP2, as shown in this study. SHIP2 is a known binding partner of Mena, and this binding can regulate its phosphatase activity ^27^. We have shown that removing SHIP2 from the cell enables further increases in ERK1/2 activity and AKT activity when Mena is overexpressed. This demonstrates that increased levels of Mena allow it to sequester SHIP2 and inhibit its phosphatase activity against ERK1/2 and AKT. While SHIP2 knockdown slightly increases ERK1/2 and AKT activity in Mena knockdown cells, it does not fully rescue phosphorylation to levels of the control, suggesting that regulation of multiple phosphatases may be involved.

### Mena as an alternative target for KRAS inhibitor-resistant PDAC

Although clinically promising, the most significant challenge to KRAS inhibitors is acquired resistance ^9–11^. We have uncovered Mena as a key regulator of KRAS^G12D^ and WT KRAS, particularly in enhancing its tumor growth and progression signaling. Our Mena knockdown data strongly support that an inhibitor against Mena would deplete signaling *via* KRAS^G12D^ and WT KRAS, decreasing tumor growth and metastasis as a result. Further studies are needed to understand the efficacy of KRAS inhibition in combination with Mena inhibition. Whether Mena inhibitors can overcome acquired resistance to KRAS inhibitors must also be explored.

### Mena as a therapeutic target for KRAS^mutant^ locoregional and metastatic PDAC

Our findings demonstrate that Mena plays a crucial role in both primary and metastatic tumor growth. As our Mena knockdown cell line was ∼85% efficient, a small population of tumor cells with a basal level of Mena expression were co-injected. After a significant period of latency, primary tumors resulted in these mice from the population of tumor cells which expressed Mena, indicating the importance of Mena for tumor initiation and growth. Furthermore, metastatic foci which spontaneously grew in these mice grew from tumor cells which expressed Mena. This supports a parallel role for Mena in tumor initiation and growth at metastatic sites and the primary tumor. Additionally, mice injected with Mena overexpressing cells produced primary and metastatic tumors which maintained robust expression of Mena, further supporting the need for Mena expression in KRAS^G12D^ primary and metastatic tumors.

Mice with Mena overexpressing tumors produced robust metastases, more frequently and larger than controls. Conversely, mice with Mena knockdown tumors produced small, infrequent metastases. Our experimental metastasis model demonstrated that this effect is due to Mena’s role in metastatic growth, rather than changes to dissemination. As Mena’s role in tumor cell dissemination is well-known and we have identified that Mena promotes PDAC tumor cell migration and invasion, it is likely that when Mena is overexpressed, increased dissemination also promotes frequent and large metastases, however the effect on metastatic growth is greater.

Overwhelming metastatic burden is the primary cause of death for patients with PDAC. This occurs due to tumor cell dissemination and rapid growth of metastases. Mena is not only a driver of tumor cell dissemination, but we have now shown that Mena plays a crucial role in promoting primary and metastatic tumor growth. Mena knockdown cells are unable to grow into primary or metastatic tumors. These findings strongly rationalize the development of a Mena inhibitor to act simultaneously as an anti-growth and anti-dissemination agent in locoregional and metastatic PDAC, as Mena is a common target in primary and metastatic tumors. Currently, no targeted therapeutics exist for metastatic PDAC and chemotherapy is used to manage metastasis growth until tumor growth outpaces treatment.

Interestingly, in our patient cohort, Mena expression levels are not significantly different, but modestly increased, in patients who had undergone neoadjuvant chemotherapy compared to those who were treatment naive (Figure S1C). Mena expression has previously been shown to increase in breast cancer patients who had undergone neoadjuvant chemotherapy compared to treatment naïve patients^44^. Mena expression is required for TMEM doorway assembly and function which is critical in the systemic dissemination of tumor cells^44^. Both considerations further rationalize the development of a Mena inhibitor as Mena’s pro-tumor effects are not mitigated following chemotherapy.

## AUTHORS’ CONTRIBUTIONS

**L.A. Ariyan**: Conceptualization, data curation, validation, visualization, investigation, methodology, analysis, writing, editing. **E.P. Zambalde**: data curation, investigation, visualization, methodology. **J. Li**: validation, methodology. **T. McAuliffe**: data curation. **E. Ma**: data curation. **F.P. Martinez**: data curation. **N. Panarelli**: data curation, analysis. **R. Eddy**: data curation, methodology. **P. Patil**: data curation, investigation, analysis. **J. Condeelis**: Conceptualization, funding acquisition, resources, methodology, review and editing. **H. Gil-Henn**: Conceptualization, investigation, methodology, supervision, visualization, review and editing. **J.C. McAuliffe**: Conceptualization, resources, investigation, data curation, supervision, funding acquisition, methodology, visualization, project administration, review and editing.

## ACKNOWLEDGMENTS

The authors are grateful for funding from the Integrated Imaging Program for Cancer Research (IIP-CR), Evelyn Gruss Lipper Foundation, and Gilead Research Scholars. Thank you to the Einstein SIG grant (1S10OD 034192-01A1). Thank you for technical and conceptual support from Camille Duran, Madeline DeLuca, Dianne Cox, Maja Oktay, David Entenberg, Analytical Imaging Facility (Vera DesMarais, Hillary Guzik,

Rotem Alon), Histology and Comparative Pathology Core, and Translational Pathology Laboratory (Aliona Zintiridou, Mary Chen).

