## Supplementary Tables and Figures for "Mena (ENAH) Promotes KRAS-Driven Tumor Growth and Metastatic Progression in Pancreatic Ductal Adenocarcinoma"

**SUPPLEMENTARY DATA**

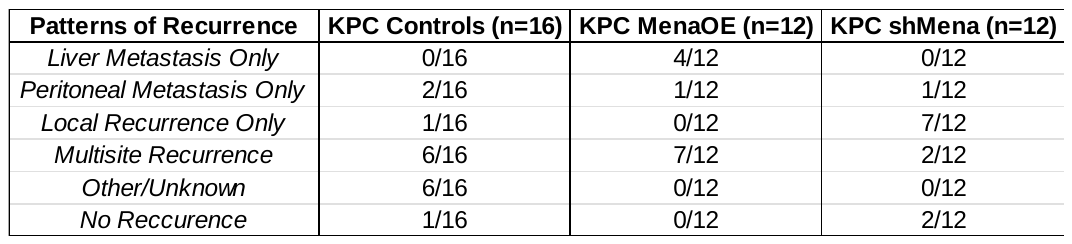

**Table S1. Patterns of Recurrence.** Recurrence type(s) were assessed post-mortem at necropsy or via H&E staining for mice who had previously undergone curative-intent primary tumor resection.

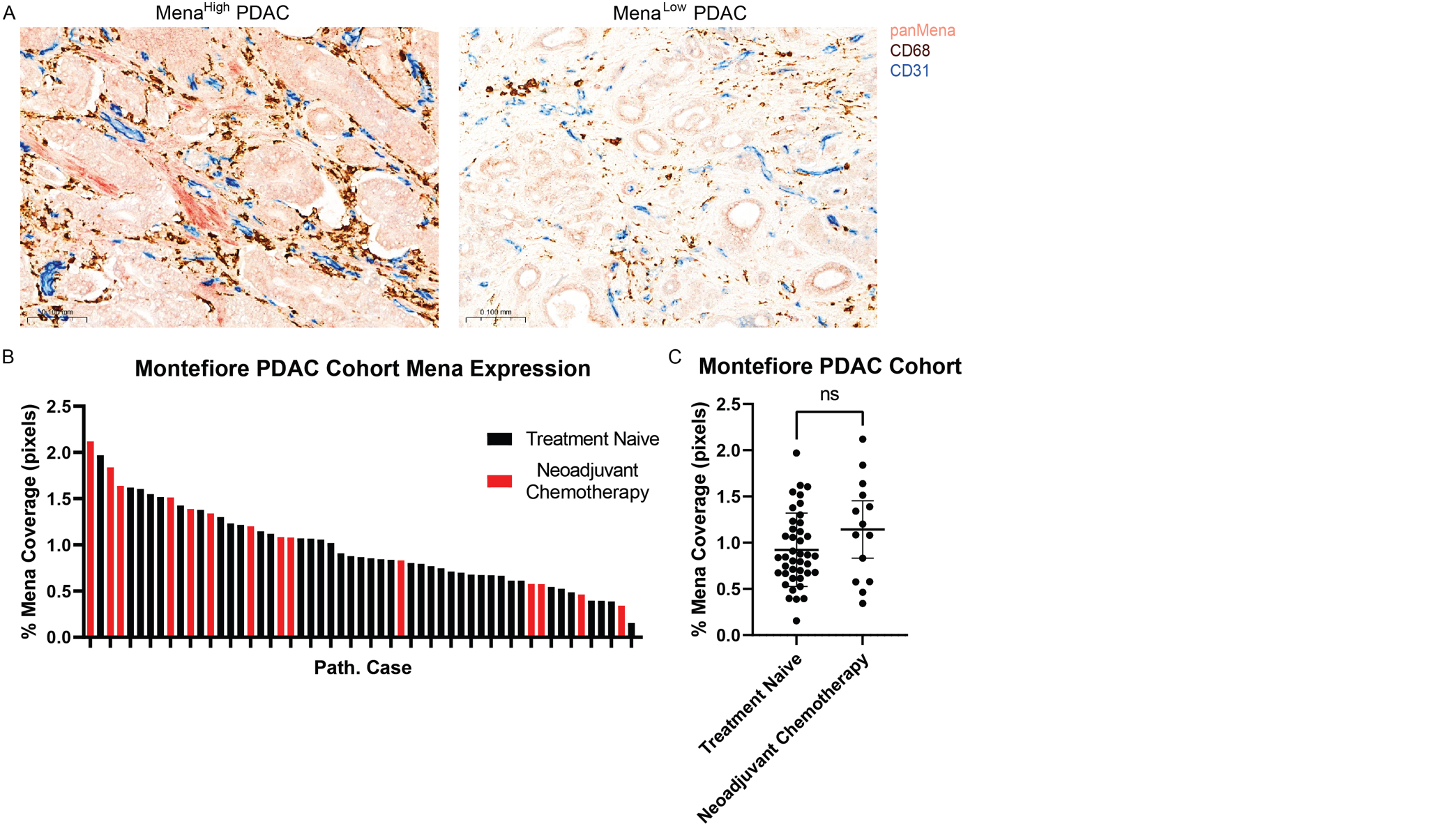

**Figure S1. Mena expression is heterogeneous in Montefiore PDAC patient cohort.** **A.** Representative images of Mena^High^ and Mena^Low^ PDAC in the Montefiore patient cohort. Validated immunohistochemical multiplex for TMEM doorways identifies Mena using red chromogen ^19,20^. **B.** Percent Mena coverage within scanned whole tissue slides shows variable Mena expression levels within the patient cohort. **C.** Percent Mena coverage is not significantly changed in patients who had undergone neoadjuvant chemotherapy (p=0.1770, Welch’s t-test).

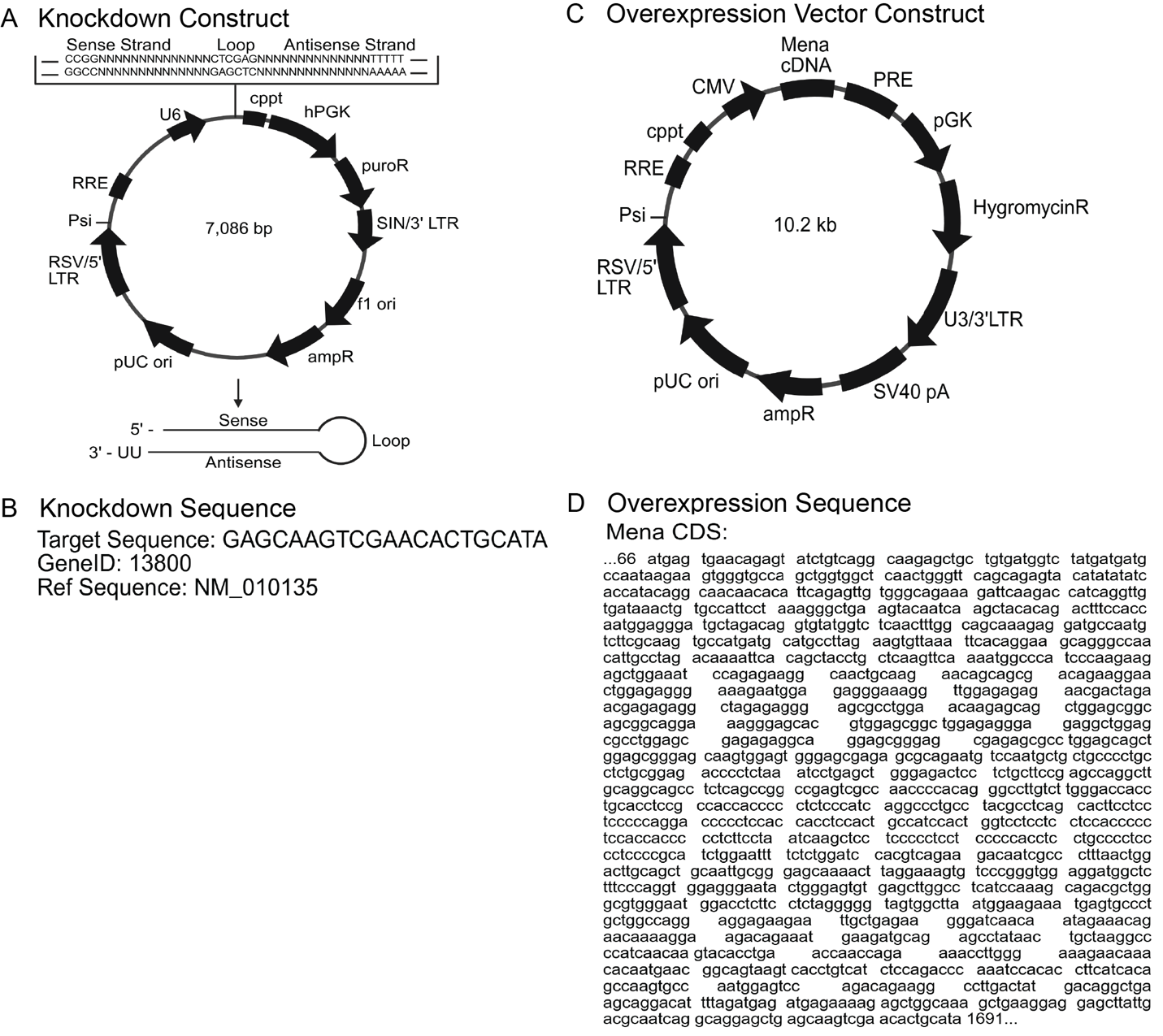

**Figure S2. Knockdown and Overexpression constructs.** **A.** Knockdown shRNA construct as outlined per manufacturer (Sigma Aldrich). Including the shRNA insert, the plasmid is 7,086bp; empty vector control plasmid is 7,052bp. The following features are included in the vector: central polypurine tract (cppt), human phosphoglycertate kinase eukaryotic promoter (hPGK), puromycin resistance gene for mammalian selection (puroR), 3’ self-inactivating long terminal repeat (SIN/3’ LTR), f1 origin of replication (f1 ori), ampicillin resistance gene for bacterial selection (ampR), pUC origin of replication (pUC ori), RSV 5’ long terminal repeat (RSV 5’LTR), RNA packaging signal (Psi), Rev response element (RRE). **B.** Selected knockdown targeting sequence and GeneID. **C.** Overexpression vector construct as modified from original construct (pLenti CMV GFP Hygro (656-4) from Eric Campeau & Paul Kaufman; Addgene plasmid # 17446). The following features are included: cytomegalovirus promoter (CMV), woodchuck post-transcriptional element (PRE), murine phosphoglycerate kinase promoter (pGK), hygromycin resistance gene (HygromycinR), U3/3’ LTR, SV40 polyadenylation signal (SV40 pA), ampicillin resistance gene (ampR), pUC origin of replication (pUC ori), RSV 5’ long terminal repeat (RSV 5’LTR), HIV-1 psi packaging signal (Psi), HIV-1 Rev response element (RRE), central polypurine tract (cppt). **D.** Inserted Mena cDNA coding sequence per Genbank.

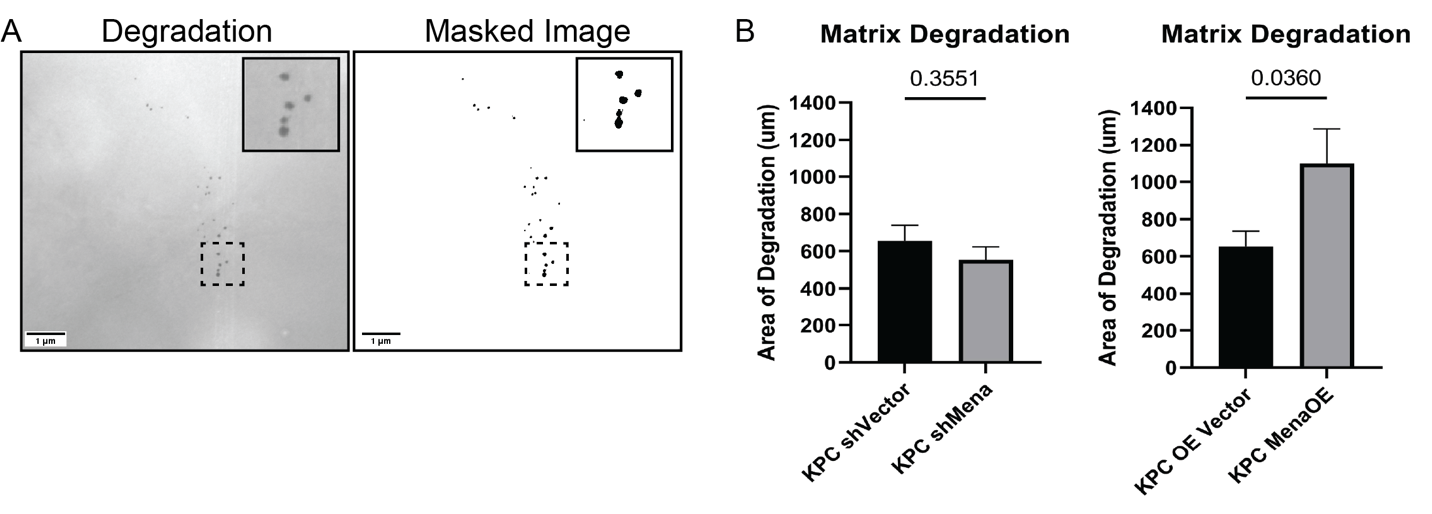

**Figure S3. In vitro matrix degradation. A.** Representative images of fluorescent matrix degradation holes (left panel) and masked image (right panel) following tumor cell incubation overnight. **B.** Plotted quantification of degradation using custom ImageJ macro; KPC shMena vs KPC shVector p=0.3551, KPC Mena Overexpression vs KPC Overexpression Vector p=0.0360.

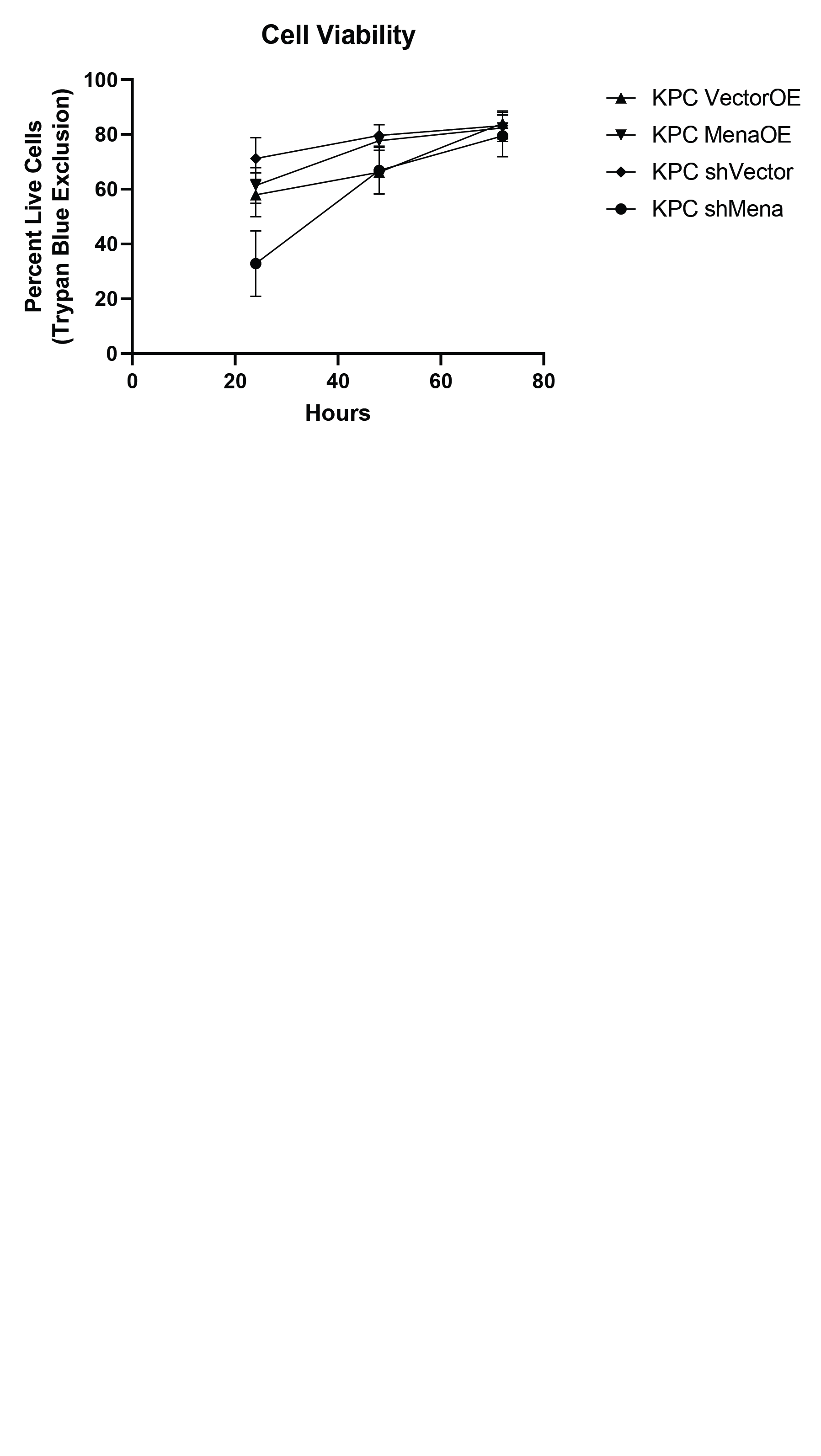

**Figure S4. In vitro cell viability.** Viability did not significantly change between controls and Mena modified cells over the course of 72 hours. KPC Mena Overexpression vs KPC Overexpression Vector, p=0.6595; KPC shMena vs KPC shVector, p=0.2904. Two-way ANOVA testing was conducted to account for changes over time between groups.

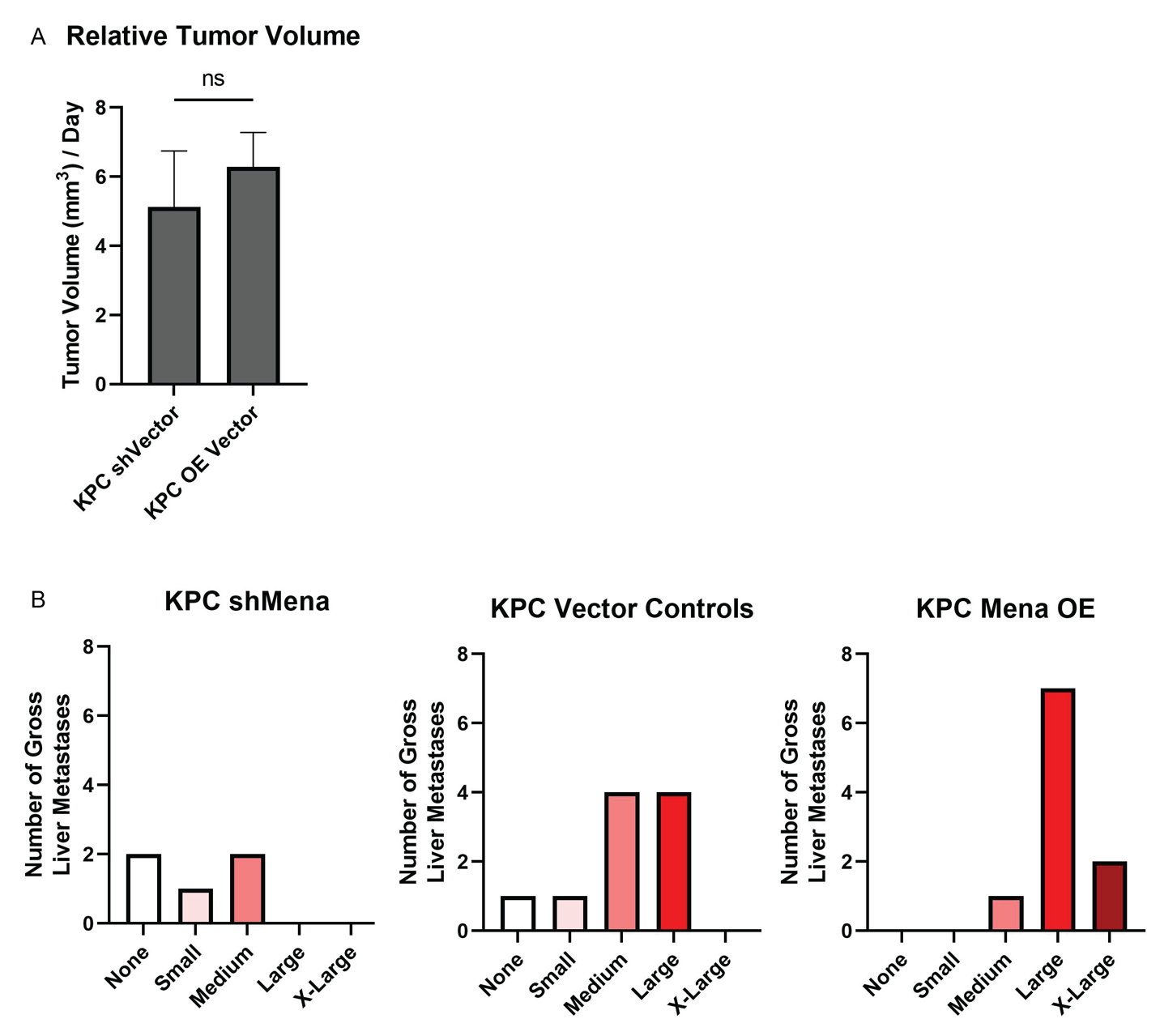

**Figure S5. Primary and metastatic tumor growth is regulated by Mena. A.** Primary tumor volume per day between vector controls. Tumor volume was measured as (length)x(width)^2^/2; KPC shVector vs KPC OE Vector, ns. **B.** Average number and size of gross liver metastases by H&E following orthotopic injection of KPC shMena (left panel), KPC vector controls (middle panel), and KPC MenaOE (right panel).

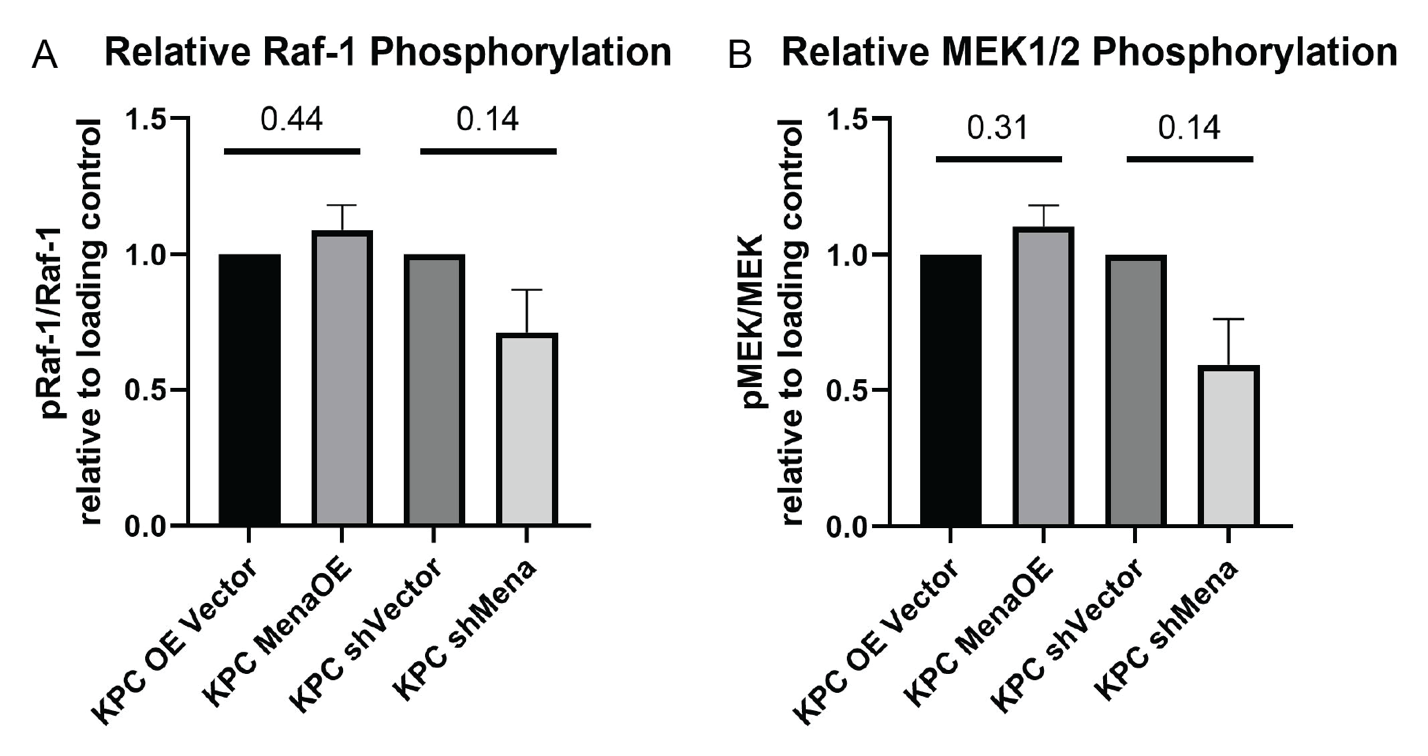

**Figure S6. Raf-1 and MEK1/2 activity are not regulated by Mena.** Phosphorylated protein was measured relative to total protein. Phosphorylated protein and total protein were both quantified relative to loading control. KPC Mena OE and KPC shMena relative protein levels were compared to normalized control vector cell lines, respectively. **A.** Phosphorylated Raf-1 does not significantly change with altered Mena levels (KPC Mena Overexpression vs KPC Overexpression Vector, p=0.44; KPC shMena vs KPC shVector, p=0.14; top panel). **B.** Phosphorylated MEK1/2 does not significantly change with altered Mena levels (KPC Mena Overexpression vs KPC Overexpression Vector, p=0.31; KPC shMena vs KPC shVector, p=0.14; center panel).

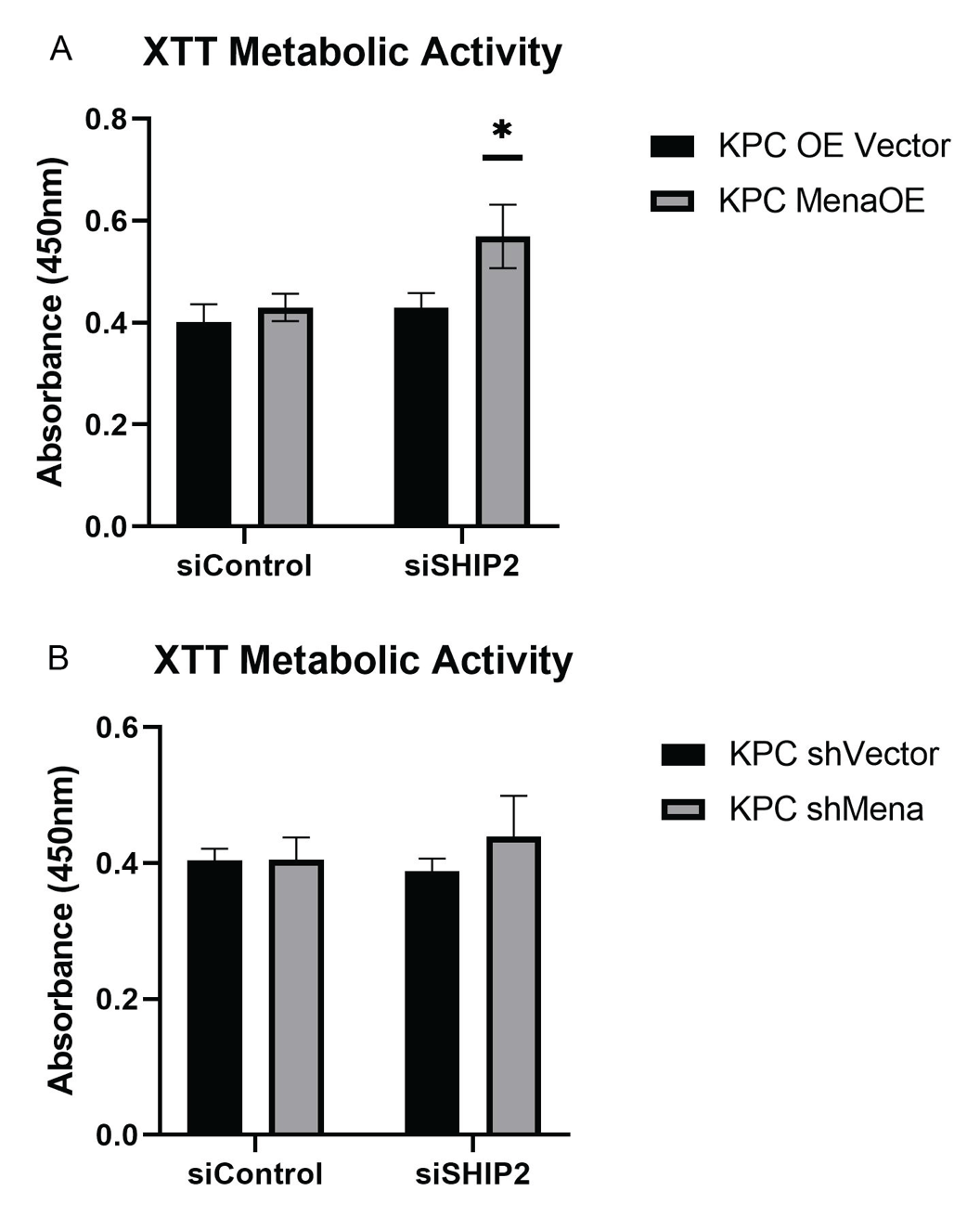

**Figure S7. XTT cell proliferation following SHIP2 knockdown.** Cell proliferation based on metabolic activity 72 hours post siSHIP2 nucleofection. A. Metabolic activity significantly increased in KPC MenaOE siSHIP2 cells (KPC OE Vector siControl vs KPC MenaOE siSHIP2, p = 0.047). B. Metabolic activity does not significantly change in KPC shMena siSHIP2 cells (KPC shVector siControl vs KPC shMena siSHIP2, p = 0.477). Statistical testing performed with Two-way ANOVA.
